# The Reissner Fiber is Highly Dynamic in vivo and Controls Morphogenesis of the Spine

**DOI:** 10.1101/847301

**Authors:** Benjamin Troutwine, Paul Gontarz, Ryoko Minowa, Adrian Monstad-Rios, Mia J. Konjikusic, Diane S. Sepich, Ronald Y. Kwon, Lilianna Solnica-Krezel, Ryan S. Gray

## Abstract

Spine morphogenesis requires the integration of multiple musculoskeletal tissues with the nervous system. Cerebrospinal fluid (CSF) physiology is important for development and homeostasis of the central nervous system and its disruption has been linked to scoliosis in zebrafish [1, 2]. Suspended in the CSF is an enigmatic glycoprotein thread called the Reissner fiber, which is secreted from the subcomissural organ (SCO) in the brain and extends caudally through the central canal to where it terminates at the base of the spinal cord. In zebrafish, *scospondin* null mutants are unable to assemble the Reissner fiber and fail to extend a straight body axis during embryonic development [3]. Here, we describe zebrafish hypomorphic missense alleles, which assemble the Reissner fiber and straighten the body axis during early embryonic development, yet progressively lose the fiber, concomitant with the emergence of body curvature, alterations in neuronal gene expression, and scoliosis in adults. Using an endogenously tagged *scospondin-GFP* zebrafish knock-in line, we directly visualized Reissner fiber dynamics during the normal development and during the progression of scoliosis, and demonstrate that the Reissner fiber is critical for the morphogenesis of the spine. Our study establishes a framework for future investigations of mechanistic roles of the Reissner fiber including its dynamic properties, molecular interactions, and how these processes are involved in the regulation of spine morphogenesis and scoliosis.

**Highlights:**

- Hypomorphic mutations in zebrafish *scospondin* result in progressive scoliosis
- The disassembly of the Reissner fiber in *scospondin* hypomorphic mutants results in the strong upregulation of neuronal receptors and synaptic transport components
- An endogenous fluorescent knock-in allele of *scospondin* reveals dynamic properties of the Reissner fiber during zebrafish development
- Loss of the Reissner fiber during larval development is a common feature of zebrafish scoliosis models

## Results and Discussion

Adolescent idiopathic scoliosis (AIS) is a complex and common disorder causing curvature of the spine. Despite accumulating evidence pointing to its heritable nature, the underlying causes are not well established. Recent efforts showed that AIS-associated variants in the centrosomal protein gene *POC5* lead to cilia defects in cell culture, and generate spine deformities when expressed in zebrafish [4, 5]. Moreover, defects in motile ependymal cell cilia leading to disruption of cerebrospinal fluid (CSF) physiology in zebrafish is also associated with scoliosis, resembling AIS [1, 2]. However, the mechanisms linking cilia and CSF defects to the onset of spine curvature in these models remain unresolved. The Reissner fiber is an enigmatic glycoprotein thread suspended in CSF of the brain and central canal of the spinal cord [6]. The chief component of the fiber is Scospondin glycoprotein, which is secreted from the sub-commissural organ (SCO) of the brain and accumulates as a structured polymer or thread [7]. Zebrafish *scospondin* null mutants produce no detectable Reissner material, fail to assemble a Reissner fiber, and display a ventral curled-down phenotype, due to a failure in straightening the body axis during embryonic development [3]. Similar defects in morphogenesis of the body axis during embryonic development have long been observed in zebrafish mutants, impaired in genes important for ciliogenesis or cilia motility [8–10] and many of these are observed with defects in Reissner fiber assembly [3]. Despite the clear contribution of CSF physiology and the Reissner fiber in body axis morphogenesis during embryonic development, it is not clear how disruptions of these processes are regulating spine morphogenesis during larval development and in the adults.

After a forward genetic screen for adult-viable scoliosis mutant zebrafish (Gray, R.S. and Solnica-Krezel, L. unpublished), whole genome sequencing, mapping, and variant calling of two non-complementing scoliosis mutations (Figure S1H, I), we identified two recessive missense alleles of the *scospondin* gene, *scospondin^stl297^* and *scospondin^stl300^*, which both disrupt evolutionarily conserved cysteine residues at independent regions of the protein (Figure S1J, K). For each allele, incross of heterozygous carriers produced morphologically normal embryos. However, at approximately two weeks postfertilization, ∼25% of the larvae displayed scoliosis in the dorsal-ventral (Figure 1B and S1B, D) and medial-lateral (Figure 1B’ and S1B’) axes, without obvious vertebral malformation. These dysmorphologies and their developmental onset resemble spine pathologies observed in adolescent idiopathic scoliosis in humans.

MicroCT analysis in developmentally-matched adult zebrafish at 90dpf revealed that *scospondin^stl297^* and *scospondin^stl300^* mutants exhibited similarities in 3D spine curvature, as well as abnormalities in bone morphology and mineralization (Figure S2). On average, patterns of sagittal and lateral displacements from midline were similar in both mutants, and most severe in posterior vertebrae (Figure S2B-C, B’-C’). In regard to bone phenotypes, both *scospondin^stl297^* and *scospondin^stl300^* mutants exhibited mildly increased bone mass (i.e., increased volume and thickness, Figure S2E-G, K-M and E’-G’, K’-M’) and mineralization (i.e., increased tissue mineral density, Figure S2H-J and H’-J’), perhaps as a response to increased mechanical loading of the spine after deformity. Neither mutant exhibited significant differences in centrum length (Figure S1A, A’), indicating that differences in body length were attributable to increased spinal curvature, rather than shortened or compressed vertebrae. When comparing phenotypic severity in mutant alleles (Figure S2N, O), z-scores for curvature indices were higher for *scospondin^stl297^* compared to *scospondin^stl300^* mutants, whereas z-scores for bone indices were generally lower.

**Figure 1.**
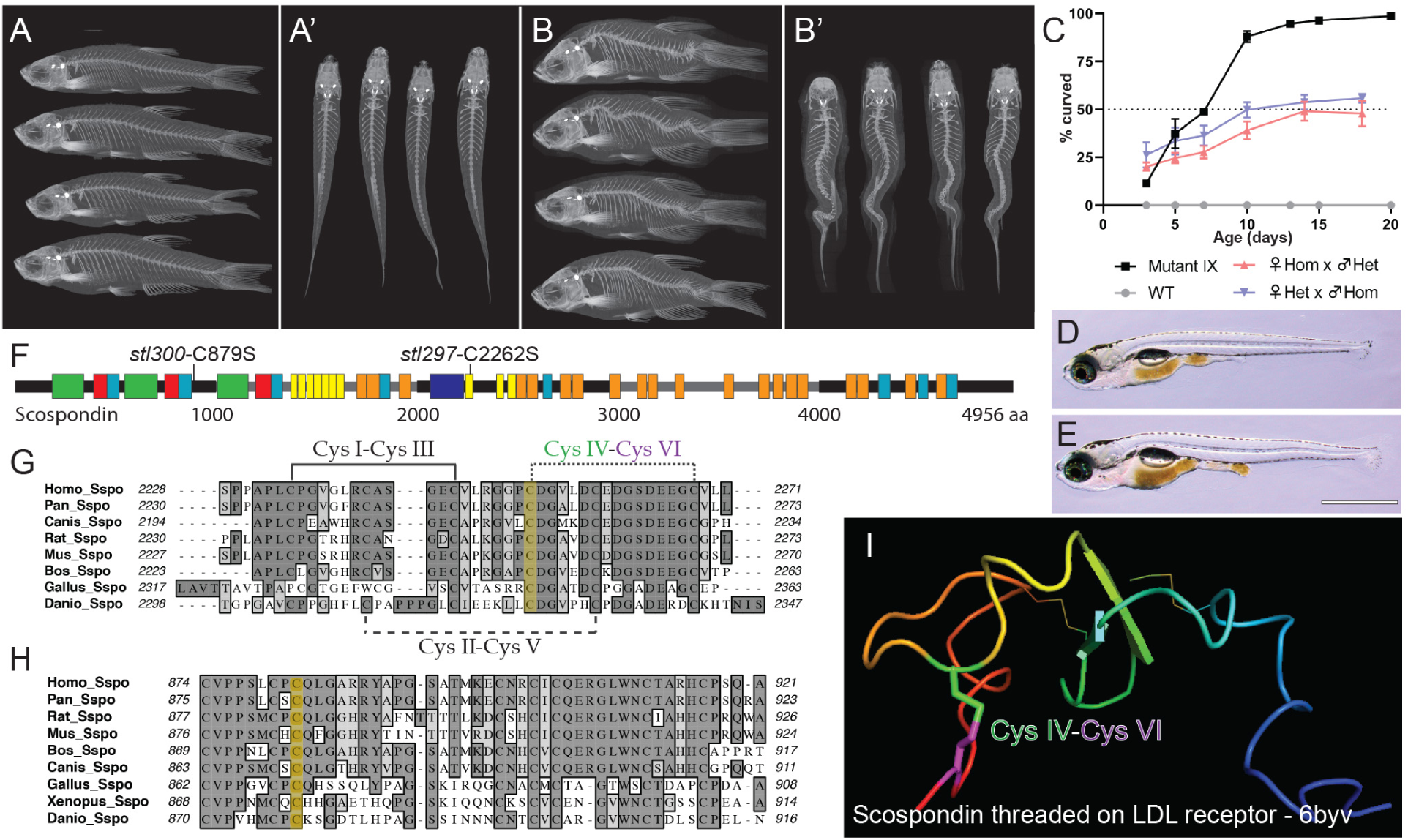
Hypomorphic mutations of *scospondin* lead to progressive scoliosis in zebrafish. (A-B’) MicroCT images of heterozygous *scospondin^stl300/+^* (A-A’) and *scospondin^stl300^* mutant zebrafish (B-B’) at 90 dpf, in both lateral (A, B) and dorsal (A’, B’) views showing adult-viable scoliosis. (C) Incidence of axial curvatures over developmental time for wild type, *MZscospondin^stl297^*, and progeny from heterozygous *scospondin^stl297/+^* x homozygous *scospondin^stl297^* mutant crosses, from both male and female homozygous *scospondin^stl297^* mutants (mean + SD; n=150, 150, 132, and 123 embryos, respectively, from three independent clutches). (D, E) Bright field image of the typical straight body of a wild-type (D) and atypical axial curvatures observed in homozygous *scospondin^stl297^* mutant (E) larvae at 13 dpf. Scale bar is 1 mm. (F) Schematic representation of Scospondin protein demonstrating the location of the *scospondin* alleles causing with scoliosis in zebrafish. (Boxes represent conserved motifs, legend in Figure S1.) (G, H) Protein alignments (Clustal-W) of Scospondin protein showing the sequence surrounding the amino acid residues (yellow highlight) affected by the *scospondin^stl297^* (G) and the *scospondin^stl300^* (H) mutations. (I) A homology model of *Danio rerio* Scospondin threaded onto the LDLrA domain derived from an LDL receptor structure (6byv). The predicted disulfide bond and labeled cysteine (Cys) residues are highlighted in (G and H). See also Figure S1–2.

For both alleles, incross of homozygous adult scoliosis mutant zebrafish resulted in 100% maternal zygotic (*MZ*) *scospondin* mutants. Which like zygotic mutants had overtly normal morphology as embryos, but displayed body curvatures by 13 dpf and adult-viable scoliosis at 2 months (Figure 1C, E and S1D). Complementation testing of *scospondin^stl297/+^* heterozygous mutants with *scospondin^Δ16/+^* heterozygous null mutants (allele described in Rose et al., co-submission) resulted in adult-viable transheterozygous *scospondin^stl297/Δ16^* scoliosis mutants (27.1%, n=236), which phenocopied *scospondin^stl297/stl297^* and *scospondin^stl300/stl300^* mutants. Furthermore, complementation testing of *scospondin^stl297/+^* heterozygous mutants with *scospondin^stl300+^* heterozygous mutants resulted in scoliosis at 13dpf (22.7%, n=198, Figure S1E). Altogether these results confirm that *scospondin^stl297^* and *scospondin^stl300^* are independent hypomorphic mutations of *scospondin* which cause viable larval and adult scoliosis in zebrafish.

A previous report of homozygous *scospondin^icm13^* null mutants demonstrated a severe posterior ventral curve down phenotype at 3 dpf [3]. In contrast, the majority of MZ*scospondin^stl297^* mutants developed a straight body axis out to 5 dpf (Figure S3A), while a few were observed with subtle axial curves (37%; *n=150*) (Figure S3B), which increased in both incidence and severity as larval development proceeded, out to 13 dpf when the majority of larvae displayed axial curvatures (94.7%; *n=113*)(Figure 1C, E). A similar progression of body curvature was observed in ∼50% of progeny when homozygous *scospondin^stl297^* mutants were crossed to heterozygous *scospondin^stl297/+^* mutant carriers, without obvious maternal effect (Figure 1C).

Scospondin protein is a large, heavily glycosylated protein composed of several repeating, well-conserved domains, established early in phylogeny [11, 12] (Figure S1L). Confirmed *scospondin^stl297^* mutant carriers are heterozygous for a T6784A (ENSDART00000097773.4) mutation (Figure 1F and Figure S1J), predicted to alter cysteine 2262 to serine (C2262S), which is located in a conserved low-density lipoprotein (LDL) receptor domain. Homology modeling of *Danio rerio* Scospondin protein sequence maps onto a crystal structure of very low-density lipoprotein receptor (6byv)(https://swissmodel.expasy.org/). This model suggests the *scospondin^stl297^* mutation could disrupt a conserved cysteine disulfide bond (CysIV-CysVI) of the LDL receptor type A motif (Figure 1G, I), which is known to be involved in protein stability of the LDL receptor [13]. Confirmed *scospondin^stl300^* mutant carriers are heterozygous for a T2635A mutation (Figure 1F and Figure S1K), predicted to alter cysteine 879 to serine (C879S), which is a highly-conserved cystiene adjacent to a trypsin inhibitor like cysteine-rich domain (Figure 1H). Homology modeling for this residue was unsuccessful.

In order to investigate the Reissner fiber in *scospondin* mutant zebrafish, we utilized an established antiserum raised against bovine Reissner fiber (AFRU) [14] for immunofluorescence. In contrast to observations in *scospondin* null mutants which fail to form a Reissner fiber at 3 dpf [3], we observed no defects in the assembly of the Reissner fiber in heterozygous *scospondin^stl297/+^* (Figure 2A) or MZ*scospondin^stl297^* mutant embryos (Figure 2B). At 5 dpf, we observed AFRU-labeled Reissner fiber and expression in the floor plate and terminal ampulla region at the base of the spinal cord in all heterozygous *scospondin^297/+^* mutant larvae (100%, n=7) (Figure 2C,E). Similarly, many of the MZ*scospondin^stl297^* larvae were observed with identical AFRU-labeled expression patterns (58%; n=19)(Figure 2D). However, several of these mutants were observed without an intact Reissner fiber or in some cases displayed only a bolus of AFRU-stained material in the central canal (42%; n=19)(Figure 2F). The presence of an intact Reissner fiber was directly correlated with a straight body axis (100%, n=11). By contrast, the absence of the Reissner fiber in these mutants was directly correlated with the onset of subtle axial curvatures in these MZ*scospondin^stl297^* mutants (100%; n=8). At 10 dpf, all *scospondin^297/+^* heterozygous mutant larvae (100%, n=7) displayed a straight body axis with an intact Reissner fiber (Figure 2G), while all the MZ*scospondin^stl297^* mutants displayed a complete absence of the Reissner fiber (100%, n=5) and axial curvatures (100%; n=5) (Figure 2H). At 10 dpf, we also consistently observed puncta of AFRU-stained Reissner material at the apical surface of individual floor plate cells in heterozygous *scospondin^stl297/+^* controls (Figure 2G), which was routinely accumulated at the basal surface of floor plate cells in the mutants at this stage (Figure 2H). Loss of Reissner fiber and atypical polarity of the Reissner material in the floor plate was also observed in MZ*scospondin^stl300^* at 10dpf (100%, n=6) (Figure S1F, G). We speculate that this may reveal a defect in the polarized secretion of Reissner material from the floor plate, which is a tissue that is known to assist in Reissner fiber assembly in zebrafish embryos [15]. In summary, we have shown that disruption of two independent, evolutionarily-conserved cysteine residues lead to disassembly of the Reissner fiber and scoliosis. It is tempting to speculate that these mutations lead to a cumulative failure of Scospondin/Reissner material secretion from the SCO and floor plate tissues during larval development. Altogether, these data suggest that the Reissner fiber has a continuous and instructive role in spine morphogenesis in zebrafish.

**Figure 2.**
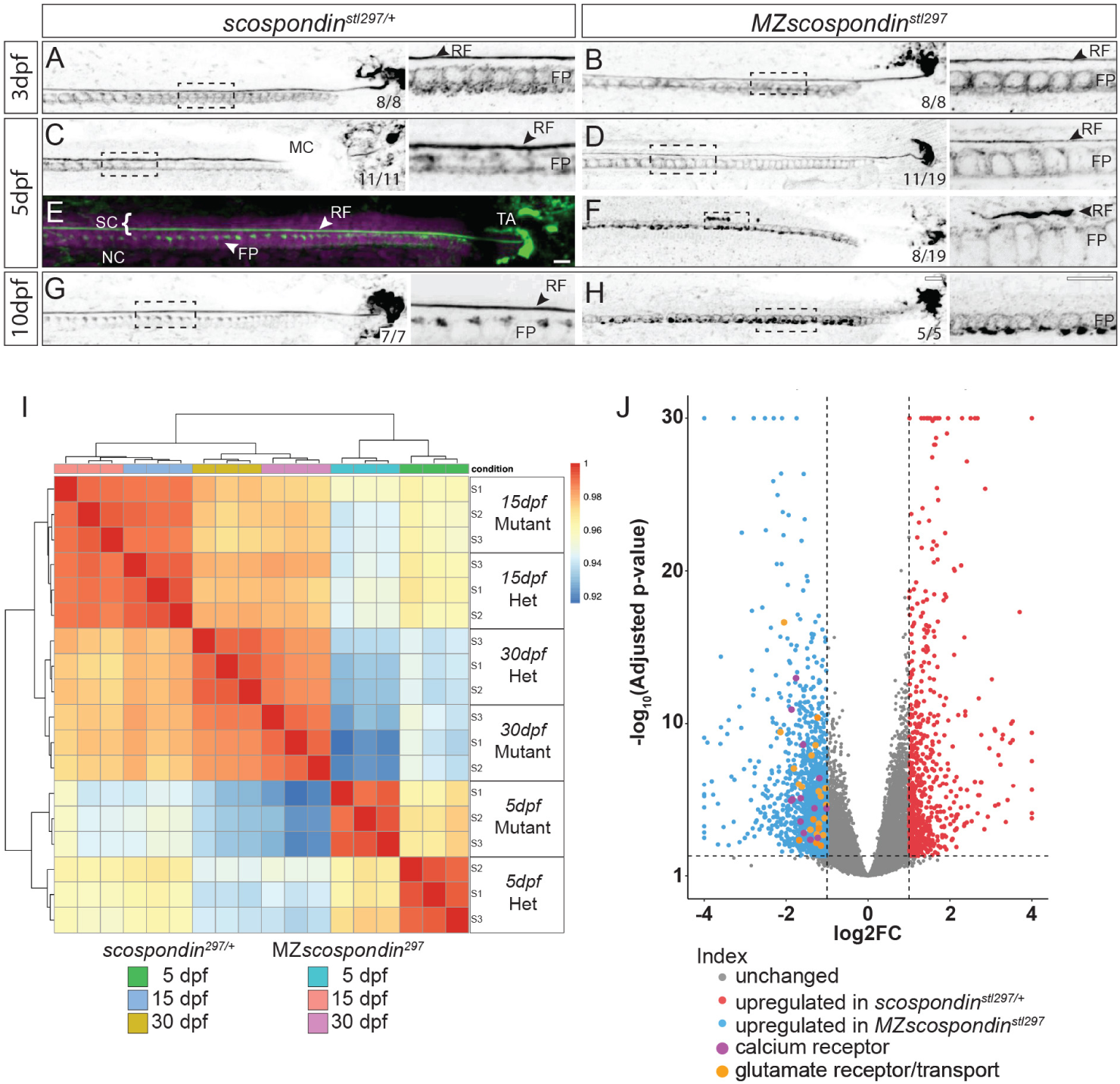
Disassembly of the Reissner fiber is correlated with the onset of axial curvatures and increased expression of several glutamate and calcium receptor genes. (A-D, F-H) Inverted greyscale maximal Z-projections of confocal stacks of the caudal region of the tail and spinal cord immunostained against the Reissner fiber (RF) in heterozygous and maternal zygotic (*MZ*) *scospondin^stl297^* mutants at 3 dpf (A, B), 5 dpf (C-F), and 10 dpf (G, H). Insets to the right of each panel to highlight a magnified region. At 3 dpf, both heterozygotes and mutants an assembled fiber (100%, n=8, 8, respectively). At 5 dpf, we observed the RF in the heterozygous mutants (100%, n=8) but the *MZ scospondin^stl297^* showed some with the fiber (58%, n=19) (D) and some with a disassembled fiber (42%, n=19)(F). Melanocytes (MC) block signal in some animals. At 10 dpf, we observed the RF in all heterozygous controls (G) which was completely lost in the mutants (H) (100%, n=7, 5, respectively). Scale bars are 10 μm. (E) Merged confocal stack showing the RF in the spinal cord (SC) (white bracket), above the notochord (NC), and terminates at the terminal ampulla (TA) with pseudocolored DAPI (magenta) and RF material (green). (I) Heatmap demonstrating the correlation of sample variation from three independent sample libraries at three distinct developmental time points from both heterozygous and maternal zygotic (*MZ*) *scospondin^stl297^* mutants. The most significant variation is observed comparing control and mutant gene expression profiles at 5dpf. (J) Volcano plot of differentially expressed genes observed in heterozygous *scospondin^stl297/+^* controls compared to *MZ scospondin^stl297^* expression at 5dpf. Upregulation of several calcium and glutamate receptors and synaptic transport components was observed in *MZ scospondin^stl297^* mutants, as labeled with larger circles in purple and orange respectively. See also Figure S3 and Table S1.

A critical question still remains of how the Reissner fiber contributes to spine morphogenesis. To address this, we sequenced bulk transcriptomes of heterozygous *scospondin^stl297/+^* and *MZscospondin^stl297^* mutants at three stages of spine development to capture: the beginning stages of Reissner fiber disassembly in early larvae (5 dpf), a late larval time-point to encompass robust axial curvature and complete loss of the Reissner fiber (15 dpf), and during skeletal maturation and robust scoliosis in the mutants (30 dpf). We generated three independent cDNA libraries for RNA sequencing at each developmental time point.

The most robust variation of differentially expressed genes amongst all samples was observed at 5 dpf (Figure 2I). Gene ontology analysis of differentially expressed genes at 5 dpf using high stringency (*p<10^-7^*) [16] revealed a strong upregulation of genes involved in transmembrane transporter activity, specifically for multiple ionotropic glutamatergic receptors, voltage-dependent calcium receptors, and a variety of related synaptic transport components in MZ*scospondin^stl297^* mutants (Figure S3C and Table S1). Analysis of the gene ontology did not reveal significant enrichment of function in downregulated genes. These data support the notion that the Scospondin-containing Reissner fiber is required for the regulation of aspects of neuronal development (*e.g.* glutamatergic neurons) in larval zebrafish, which may contribute disruption of normal neuromuscular developmental programs leading to the onset of axial curvatures through. Indeed, Scospondin has been previously shown to affect neurogenesis *in vitro* [17, 18]. The specific nature of any alterations in neuromuscular development at a cellular and physiological level in the *scospondin* hypomorphic mutant zebrafish alleles presented here, requires further investigation.

To monitor the dynamic properties of the Reissner fiber during axis straightening we engineered a fluorescent *scospondin* knock-in allele (*scospondin-GFP^ut24^*) in zebrafish. In brief, we used CRISPR/Cas9 to promote a targeted double strand break within the last exon of the *scospondin* gene and an EGFP donor cassette with homology arms for in-frame C-terminal tagging (Figure S4A). From the resulting adult F0 fish, we isolated a single founder male by (i) PCR screening for EGFP sequence in genomic DNA from isolated sperm samples; (ii) by confocal imaging of outcrossed progeny; and (iii) by co-localization with AFRU immunofluorescence (see methods). At 3 dpf, we observed Scospondin-GFP expression as an extremely straight Reissner fiber extending from head to tail (Figure 3A-B’ and S4B). In the head, we observed Scospondin-GFP expression in the most rostral SCO and in the flexural organ (Figure 3A, A’ and S4B-C’). In the tail, we observed expression in the floor plate and the Reissner fiber ending as a coiled mass within the terminal ampulla at the base of the spinal cord (Figure 3B, B’ and S4B’). We continue to observe the presence of the Scospondin-GFP labeled Reissner fiber and terminal ampulla in young adult fish (Figure S4D, D’). As expected, endogenous Scospondin-GFP expression is similar to previously reported expression patterns in zebrafish based on Reissner fiber antibodies and whole-mount *scospondin* gene expression [3, 12]. In agreement, Scospondin-GFP expression in *scospondin-GFP^ut24^*, knock-in zebrafish displayed tight colocalization (Pearson’s R value, 0.98) with the expression pattern of AFRU antiserum expression [14] (Figure 3C-E).

**Figure 3.**
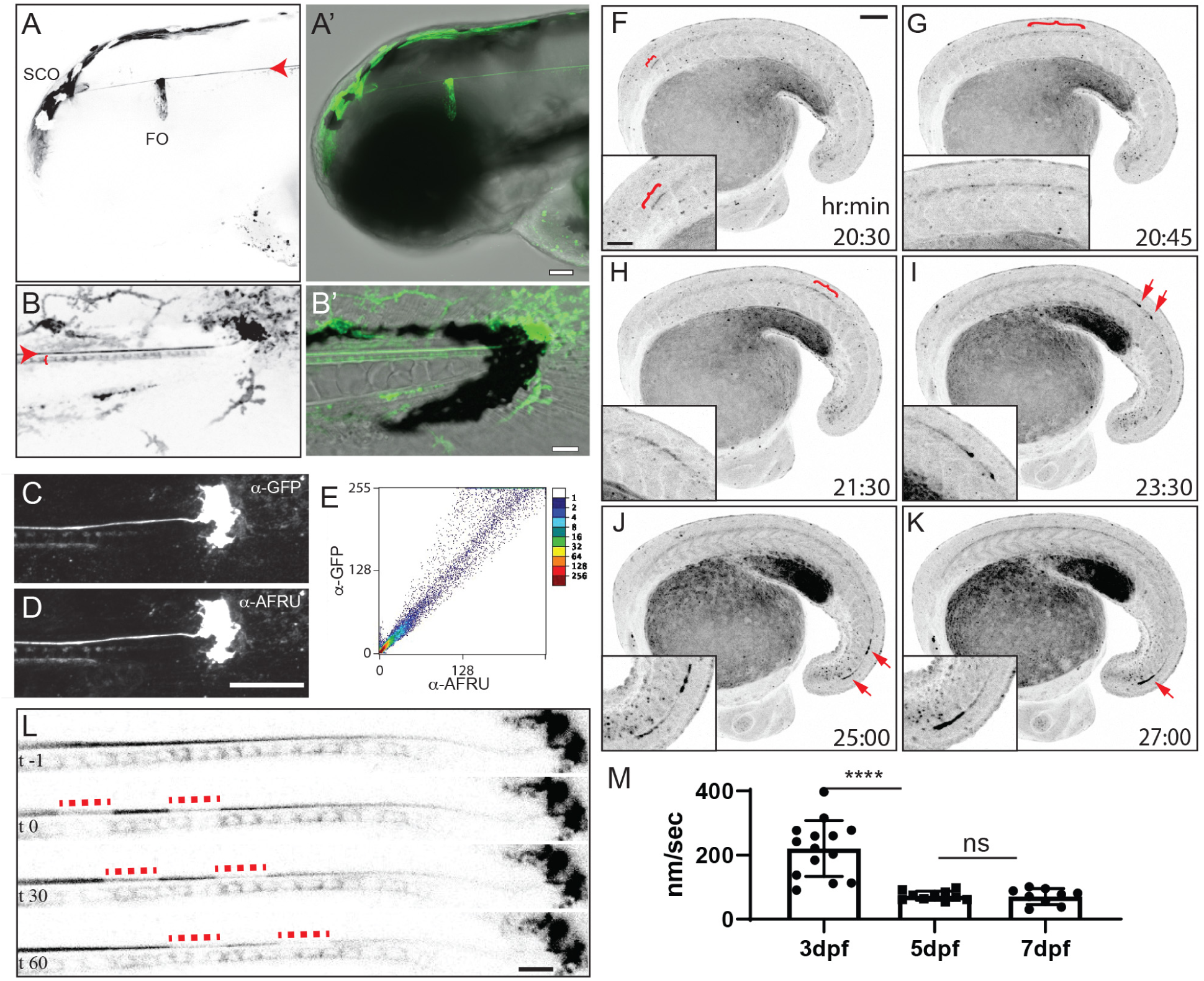
Dynamic properties of the Reissner fiber revealed during development in *scospondin-GFP^ut24^* knock-in zebrafish. (A, B) Inverted greyscale maximal Z-projections of confocal stacks of *scospondin-GFP^ut24/+^* embryos at 3 dpf. Expression in the subcommissural organ (SCO) and flexural organ (FO) in the head (A) and the Reissner fiber (RF) (red arrowhead) and floor plate in the tail (red bracket) (B). (A’, B’) merge of DIC image and pseudocolored Scospondin-GFP expression (Green) as in (A, B). Scale bars: 50 μm (A, A’) and 20 μm (B, B’). (C-E) Colocalization analysis of greyscale maximal Z-projections of confocal stacks of Scospondin-GFP expression in *scospondin-GFP^ut24/+^* knock-in zebrafish using anti-GFP (C) and AFRU antiserum (D) demonstrated a high degree of signal colocalized (Pearson’s R^2^ = 0.98). Scale bars: 25 μm. (F-K) Frames from a time-lapse confocal dataset taken during tail bud development (20-30 hours post fertilization, see Video S1) presented as inverted greyscale maximal Z-projections. Red brackets highlight faint expression, extended fibers of Scospondin-GFP, while red arrows point out a bolus Scospondin-GFP material which is observed to travel rapidly in a rostral to caudal direction. Inset in the lower left-hand side of each panel are digitally enlarged portions of the region containing Scospondin-GFP -labeled material. Scale bars: 100 μm in main, 50 μm in inset (L) Inverted greyscale maximal Z-projections of confocal imaging of a representative *scospondin-GFP^ut24/+^* knock-in embryo at 3 dpf. At time (t)=0, two regions were photobleached using a short, high energy pulse from a 488nm solid state laser which allowed for manual tracking of the movement of the bleached region from rostral to caudal. Scale bar 10μm. (M) Average velocity as nanometers (nm) per second (sec) were calculated manually for multiple embryos at 3, 5, and 7dpf (n=13, 9, and 8 respectively). The average velocity for each individual embryo was plotted as box plots (mean ± SD). (**** denotes p-value <10^-4^) See also Figure S4 and Video S1-4.

To describe the dynamics of the Reissner fiber assembly during development we used confocal time lapse imaging in *scospondin-GFP^ut24/+^* embryos during tail morphogenesis. We first observed the assembly of Scospondin-GFP-labeled fiber in the rostral spinal canal ∼20.5 hours post fertilization (hpf) (red bracket, Figure 3F and Video S1). Shortly after, we also observed a larger fiber of Scospondin-GFP extending more caudally (red bracket, Figure 3G, H). This Scospondin-GFP material was also observed to accumulate as a bolus of material (red arrows, Figure 3I, J), which was observed to rapidly travel in a rostral to caudal direction to join established Scospondin-GFP labeled fiber at the end of the spinal cord (Figure 3K). During robust axial elongation (2-3dpf), we observed a highly dynamic breakdown and distribution of Scospondin-GFP signal outward from the terminal ampulla and into the developing fin fold (Video S2). It has long been suspected that the Reissner fiber continually grows in a rostral-caudal direction [19], consistent with the observations of directional transport of labeled-monoamines along the fiber in rat [20]. To quantify fiber motility during zebrafish development, we photobleached Scospondin-GFP-labeled Reissner fiber to mark regions for tracking (Figure 3L and Video S3). At various stages of development, we observed that photobleached regions of the Reissner fiber always traveled in a continuous rostral to caudal direction. The average speed at 3 dpf was 220±87nm/sec. In contrast, at 5 and 7 dpf the speed of the fiber was significantly slower (70±14nm/sec and 70±24nm/sec, respectively; t-test; *p*=1.37×10^-5^; Figure 3L, M). Using high-speed imaging (10Hz) at higher magnification we also observed rapid movement of Scospondin-GFP-labeled puncta along the Reissner fiber in a rostro-caudal direction, which were occasionally observed to extend away from the fiber, towards the floor plate, and retract back into the bulk Reissner fiber (Figure S4E, Video S4). In summary, we have described several dynamic properties of Reissner fiber during early zebrafish development by endogenous fluorescent Scospondin-GFP expression. This knock-in strain and observations set the stage for future studies aimed at defining molecular interactions of the Reissner fiber and the mechanisms driving the rostro-caudal motility of the fiber.

As *scospondin-GFP^ut24^* is an endogenous knock-in into a wild-type allele, we are currently precluded from dynamic imaging of the Reissner fiber in the hypomorphic *scospondin* mutants reported here, using this approach. However, there are obvious phenotypic similarities between *scospondin* hypomorphic mutants and other cilia-related scoliosis mutants described previously [1, 2]. We hypothesized that the loss of the Reissner fiber is a common phenotype associated with the onset of scoliosis in zebrafish. To test this, we crossed *scospondin-GFP^ut24^* to a dominant enhancer-trap transgenic scoliosis mutant, *Et(druk-GFP^dut26/+^)*. This mutant was generated by a fortuitous, Tol2-GFP integration which displays a unique GFP-expression pattern which is tightly linked with the onset of adult-viable scoliosis (98%; *n=981*), without vertebral malformation (Gray, R.S., McAdow, A.R., Solnica-Krezel, L. and Johnson, S.L., unpublished). At both 3 and 5 dpf, all *Et(druk-GFP^dut26/+^);scospondin-GFP^ut24/+^* double mutant, knock-in larvae displayed Scospondin-GFP-labeled Reissner fiber (Figure 4A-B’). However, at the onset of scoliosis in *Et(druk-GFP^dut26/+^);scospondin-GFP^ut24/+^* mutants, we observed a consistent loss of Reissner fiber (100%; *n=9*)(Figure 4D, D’). Mutations in the *kinesin family member 6* (*kif6*) gene cause a loss of ependymal cell cilia and scoliosis, without vertebral malformations in zebrafish [1, 21]. We assayed the Reissner fiber using AFRU immunostaining in *kif6^sko/sko^* mutants which begin to display subtle axial curvatures at 3dpf [21]. Interestingly at this time, we also observed strong defects of Reissner fiber assembly (Figure 4F), with only occasional tangles (Figure 4F’) or puncta (Figure 4F’’) of Reissner material present. Altogether the loss of Reissner fiber in two independent models of scoliosis confirms it role in the homeostasis of the straight body axis and spine morphogenesis in zebrafish.

**Figure 4.**
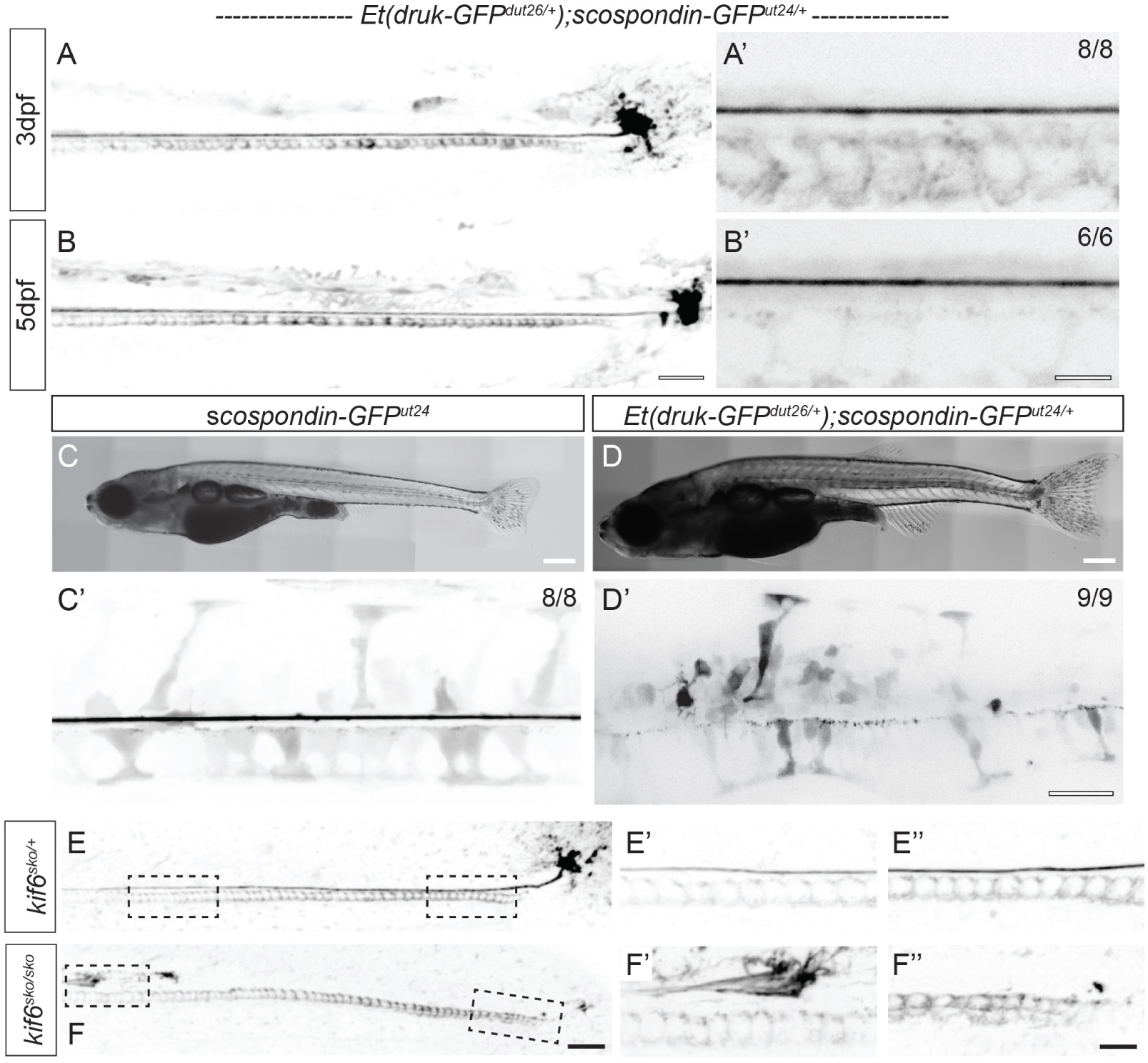
Loss of the Reissner fiber is a common phenotype associated with scoliosis in zebrafish. (A-B’) Inverted greyscale maximal Z-projections of confocal imaging of *Et(druk-GFP^dut26/+^); scospondin-GFP^ut24/+^* demonstrating typical assembly of the Reissner fiber and floor plate and terminal ampulla expression in these mutant knock-in embryos at 3dpf (A, A’) and 5dpf (B, B’). Scale bars are 25μm in (A, B) and 5μm in (A’, B’). (C, D) Stitched bright field images of a wild-type juvenile *scospondin-GFP^ut24/+^* (C) and an aged-matched *Et(druk-GFP^dut26/+^; scospondin-GFP^ut24/+^* displaying the onset of mild scoliosis. (C’, D’) Inverted greyscale maximal Z-projections of confocal imaging of the same fish in C and D respectively to highlight the spinal cord. At 19dpf, in wild-type *scospondin-GFP^ut24/+^* we observe high expression of the Reissner fiber (100%, n=8)(C’), while in scoliosis mutant *Et(druk-GFP^dut26/+^; scospondin-GFP^ut24/+^* at 19dpf we observed curvature of the spinal canal and a complete loss of a Reissner fiber (100%, n=9). Scale bars are 500μm in (C, D) and 25μm in (C’, D’). (E-F’) Inverted greyscale maximal Z-projections of confocal imaging of AFRU-stained zebrafish embryos at 3dpf. Normal Reissner fiber is observed in heterozygous kinesin family member 6 (*kif6*) mutants at multiple axial levels (8/8, E-E’’). In contrast, homozygous *kif6^sko^* mutants show severe disassembly of the Reissner fiber, with occasional tangles (F’) or puncta (F’’) of Reissner material in the central canal (7/8).

Since the discovery of the Reissner fiber multiple hypotheses have been proposed for its function including, detoxification and transport of molecules in CSF [20], neurogenesis during early brain development [17, 18], and through its direct interaction with the ciliated CSF-contacting neurons lining the central canal, as a mechanosensory organ controlling the “flexure of the body” [22, 23]. Moreover, earlier work in amphibians demonstrated that the resection of the SCO disrupted Reissner fiber assembly, led to scoliosis in some animals [24, 25]. Here, we demonstrated that two independent, evolutionally-conserved cysteine residues in Scospondin are critical for stability of the Reissner fiber during larval development. One of these cysteines (C2262) is predicted to form a disulfide bridge in one of several canonical LDL receptor A domains found in Scospondin. Interestingly, the LDL protein Apolipoprotein B has been directly visualized in the central canal in zebrafish [26] and is found in the CSF of rat and humans by proteomic analysis [27–29]. Apolipoprotein B is also an important neurogenic factor *in vitro* [30], is important for brain development in mice [31], and forms a complex with Scospondin CSF which can synergistically modulate neurodifferentiation in organotypic brain culture [32]. For these reasons, it is tempting to speculate that the C2262S mutation may also disrupt important LDL interactions with the Reissner fiber, causing alterations in neuronal differentiation in *scospondin^stl297^* mutant zebrafish. Our results using a variety of genetic manipulations now demonstrate that the intact and likely dynamic nature of the Reissner fiber *in vivo*, is required for maintaining a straight body axis and spine morphogenesis. Whereas its disassembly is driving alterations in neuronal gene expression and the onset of atypical axial curvature, which leads to progressive scoliosis in adult zebrafish. Our study opens up a new field of exploration of the mechanisms of Reissner fiber dynamics, the importance of molecular interactions of the fiber and CSF components for neuronal development, and whether these mechanisms are more generally applicable for scoliosis in humans.

## STAR METHODS

### LEAD CONTACT AND MATERIALS AVAILABILITY

Further information and requests for reagents should be directed to the lead contact, Ryan S. Gray.

### EXPERIMENTAL MODEL AND SUBJECT DETAILS

#### Zebrafish Maintenance and Care

All experiments were performed according to University of Texas at Austin IACUC standards. Wild type AB strains were used unless otherwise stated. Embryos were raised at 28.5 °C in fish water (0.15% Instant Ocean in reverse osmosis water) and then transferred to standard system water at 5dpf.

#### WGS / WES analysis

Phenotypic, mutant zebrafish were pooled and submitted for sequencing. Non-phenotypic wild type and heterozygous siblings were pooled together and submitted for sequencing. Raw reads were aligned to zebrafish genome GRCz10 using bwa mem (v0.7.12-r1034) with default parameters and were sorted and compressed into bam format using samtools (v1.6) [33]. Variants were called using bcftools (v1.9) functions mpileup, call, and filter. mpileup parameters “-q 20” and “-Q 20” were set to require alignments and base calls with 99% confidence to be used and filter parameters “-s LowQual -e ’%QUAL<20’” were used to remove low quality variant calls. Variants were then annotated and filtered by an in-house pipeline. Briefly, variants occurring with the same allele frequency between phenotypic and aphenotypic samples were filtered from further analysis as were variants that were not called as being homozygous in the phenotypic sample. Variants in the mutant samples that were homozygous for the wild type allele were also excluded. Variants found in the dbSNP database (build v142) of known variants were also filtered out and excluded from further analysis. The wild type and mutant alleles at each variants site were tabulated, and the Fisher p-value was calculated for each variant site. These remaining variants classified based on their genomic location as being noncoding site variants, coding site variants, or variants that may affect gene splicing using zebrafish Ensembl annotation build v83. For coding site variants, the amino acid of the wild type allele and the mutant allele were determined from the Ensembl annotation. The p-values were plotted against genomic location, and a region of homozygosity in the genome with a cluster of small p-values was found. Variants occurring within this region of homozygosity were manually prioritized for nonsynonymous mutations in genes.

#### MicroCT scanning and analysis

MicroCT scanning was performed as previously described [34]. All analyses were performed in precaudal and caudal vertebrae only (we refer to the 1^st^ precaudal vertebrae as vertebra 1). For analysis of spinal curvature, centrum centroid positions were identified in maximum intensity projections. A line was drawn connecting the first and last vertebrae. The absolute value of the displacements from this line were computed in the sagittal and frontal planes to compute Sagital Displacement (Sag.Disp) and Lateral Displacement (Lat.Disp) for vertebrae 1-20. For analysis of bone, FishCuT was used to quantify Length (Le), Volume (Cent.Vol), Tissue Mineral Density (TMD), and Thickness (Th) in the Centrum (Cent), Neural Arch (NA), and Haemal Arch (Haem) for vertebrae 1-16 [34]. Computation of standard scores, z-scores, and statistical testing using the global test were performed as previously described [34, 35].

#### Generation of endogenously tagged scospondin-GFP^ut24^ allele

CRISPR/Cas9 targets were chosen using CHOPCHOP and guides were synthesized according to the CHOPCHOP protocols (Montague et al., 2014; Labun et al., 2016, 2019). The last exon of *scospondin* was targeted with the CRISPR guide AGTGTACCAGCTGCCAGGGT<u>GGG</u> (PAM underlined) predicted to cut 6 bp upstream of the stop codon. To generate an sgRNA guide, an oligo containing a T7 promoter, gene-specific targeting sequence, and annealing region was synthesized (Sigma-Aldrich). This oligo was annealed to a generic CRISPR oligo using CloneAmp Hifi Polymerase. RNA was synthesized using the NEB HiScribe T7 RNA synthesis kit and purified with the Zymo RNA Clean and Concentrator-5 kit.

A plasmid was constructed to serve as a donor. The plasmid contained 5’ and 3’ homology arms (776 and 532 bp respectively) flanking the eGFP coding sequence (720bp). The donor was constructed such that the PAM would be abolished and the eGFP coding sequence would be inserted just before the endogenous stop codon. The homology arms were cloned from wild-type AB* zebrafish DNA and eGFP was cloned from the p3E-2AnlsGFP plasmid (Kwan et al., 2007) using Clontech Hifi polymerase mix. These three fragments were purified (NucleoSpin Gel and PCR Clean-up, Machery-Nagel #740609) and Gibson cloned into the EcoRI site of pCS108 using the In-Fusion HD Cloning kit (Clontech) following manufacturer protocols.

Wild type AB zebrafish were incrossed and one-cell embryos were injected with 1nL of injection mix containing 5µM EnGen Spy Cas9 NLS (NEB #M0646), 100 ng/µL sgRNA, and 25 ng/µL donor plasmid. Once the embryos reached adulthood, sperm was collected from F0 males and DNA was extracted by diluting sperm into 50uL of 50mM NaOH and heated to 95°C for 40 minutes. PCR was performed with GFP-specific primers using GoTaq Green Master Mix (Promega #M7123). Males that generate an amplicon after PCR were outcrossed to WT AB females, and the progeny were screened for GFP expression in the SCO and Reissner Fiber.

#### Skeletal preparation

Animals were euthanized in tricaine and fixed in 10% formalin overnight, then incubated in acetone overnight. Acetone was washed away with water, and animals were stained with Bone/Cartilage Stain (0.015% Alcian Blue, 0.005% Alizarin Red, 5% Acetic Acid, 59.5% Ethanol) at 37°C overnight, and cleared in 1% KOH for days to weeks depending on the size of the fish. The fish were then moved to 25%, 50%, and then 80% glycerol and imaged.

#### AFRU immunofluorescence of Reissner Fiber

Animals were euthanized with high dose tricaine (MS-222), and then fixed in ice-cold methanol overnight. Animals were washed in PBSTr (1x PBS with 0.1% Triton X-100), blocked with 10% normal goat serum in 1x PBS with 0.5% Triton X-100. AFRU primary antibody was shared by Esteban Rodriguez (Rodríguez *et al.*, 1984) and used at 1:2000 dilution in blocking solution overnight at room temperature. Specimens were then washed and stained with secondary goat anti rabbit H+L Alexa488 at 1:1000 overnight at 4°C, washed in PBSTr and counterstained with DAPI. Specimens were immobilized in 1% low-melt agarose in 1x PBS and imaged with a Nikon Ti2E with a CSU-W1 spinning disc confocal system.

#### RNA sequencing

Library preparation was performed with 10ng of total RNA, integrity was determined using an Agilent bioanalyzer. ds-cDNA was prepared using the SMARTer Ultra Low RNA kit for Illumina Sequencing (Clontech) per manufacturer’s protocol. cDNA was fragmented using a Covaris E220 sonicator using peak incident power 18, duty factor 20%, cycles/burst 50, time 120 seconds. cDNA was blunt ended, had an A base added to the 3’ ends, and then had Illumina sequencing adapters ligated to the ends. Ligated fragments were then amplified for 12 cycles using primers incorporating unique index tags. Fragments were sequenced on an Illumina NovaSeq using paired end reads extending 150 bases.

#### RNA Quantification and Statistical Analysis

Raw reads were first trimmed using cutadapt to remove low quality bases and reads [36]. Trimmed reads were then aligned to zebrafish genome GRCz10 with ensembl annotation v83 using STAR (v2.5.4) with default parameters [37]. Transcript quantification was performed using featureCounts from the subread package (v1.4.6-p4) [38]. Differential expression analysis was performed by using the R package DESeq2 in negative binomial mode using quantified transcripts from featureCounts [39]. A two-fold expression change, expression higher than 1CPM, and a Benjamini and Hochberg false discovery rate less than 0.05 were set as a cutoff for a gene to considered differentially expressed.

### DATA AVAILABILITY

RNAseq data (GSE138842) and WGS/WES sequencing data (GSE138920) generated during this study are available at the SRA browser.

## Acknowledgements

We thank Esteban Rodriguez and Maria Montserrat Guerra (Universidad Austral de Chile) for their generous gift of the AFRU antiserum. We thank Sierra Szkrybalo for assistance with zebrafish experiments. We thank Drs. John Wallingford and Claire Wyart for critical comments this manuscript. We thank the Genome Technology Access Center in the Department of Genetics at Washington University School of Medicine for help with genomic analysis. The Center is partially supported by NCI Cancer Center Support Grant #P30 CA91842 to the Siteman Cancer Center and by ICTS/CTSA Grant# UL1TR002345 from the National Center for Research Resources (NCRR), a component of the National Institutes of Health (NIH), and NIH Roadmap for Medical Research. This publication is solely the responsibility of the authors and does not necessarily represent the official view of NCRR or NIH. Research reported in this publication was supported in part by a grant from Spinal Cord Injury/Disease Research Program (SCIDRP) (LSK and Steve Johnson), the National Institute of Arthritis and Musculoskeletal and Skin Diseases of the National Institutes of Health under Award Number R01-AR072009 to (R.S.G.) and by the National Institute for Child Health and Human Development of the National Institutes of Health under Award Number P01HD084387.

---

**Figure S1.**
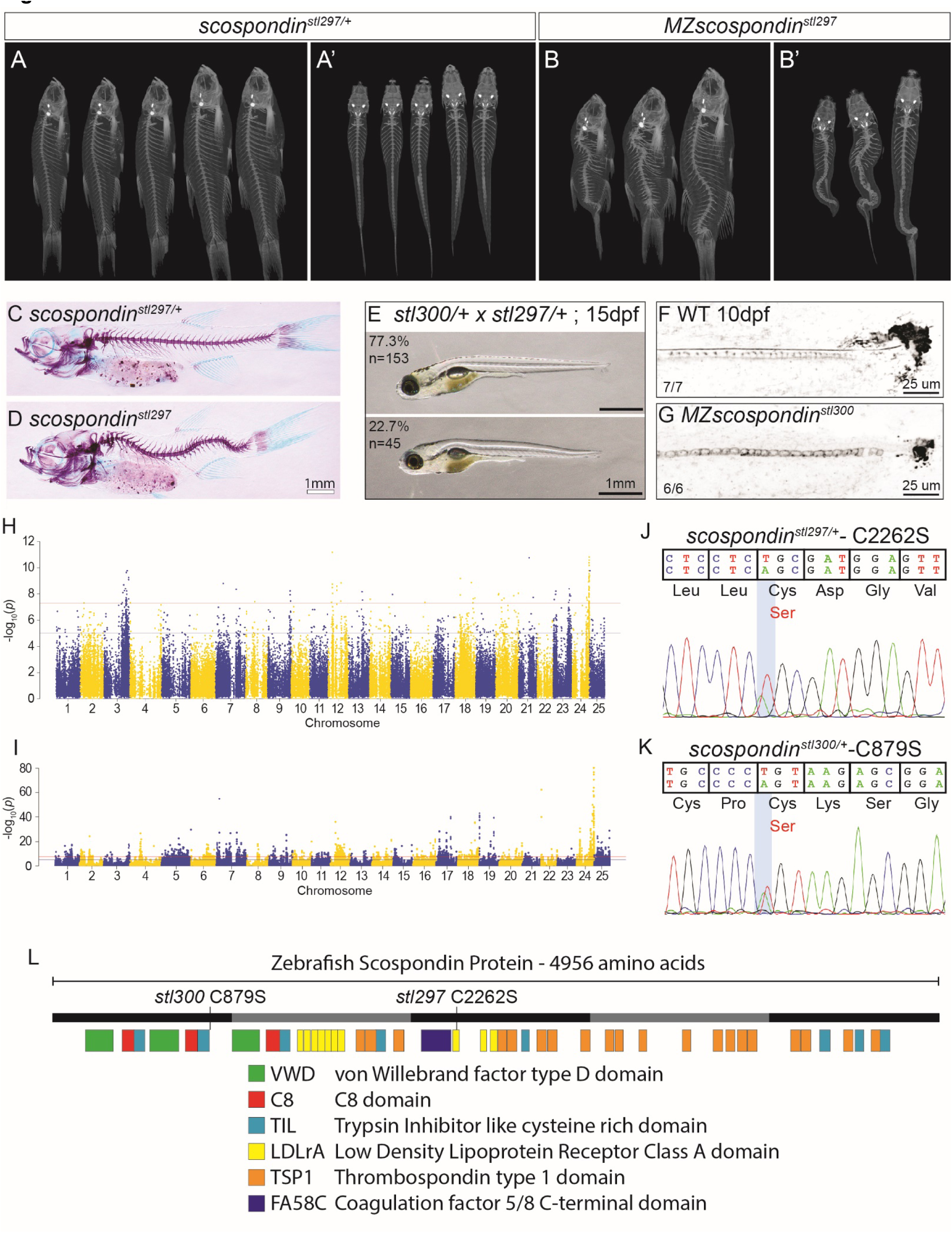
Mapping and description of *scospondin* (*scospondin*) mutant zebrafish. (A-B’) MicroCT images of heterozygous *scospondin^stl297/+^* (A-A’) and *scospondin^stl297^* mutant zebrafish (B-B’) at 90 dpf, in both lateral (A, B) and dorsal (A’, B’) views. (C, D) Alizarin-Red/Alcian-Blue skeletal staining and lateral view of heterozygous *scospondin^stl297/+^*(C) and *scospondin^stl297^* mutant (D) zebrafish at 2 months. (E) Bright field images of representative phenotypes at 15 dpf from complementation testing of a heterozygous *scospondin^stl297/+^* and *scospondin^stl300/+^* incross. (F, G) Inverted greyscale maximal Z-projections of confocal stacks of tail tips immunostained against the Reissner fiber in 10dpf wild type (F) and 10spf *MZscospondin^stl300^* mutants (G). (H, I) Manhattan plots of the ratio of mutant SNP frequency in the mutant and wild-type sibling pools using whole genome sequence from *scospondin^stl297/+^* incrosses (H) and whole exome sequence from *scospondin^stl300/+^* incrosses (I). (J, K) Sanger sequence traces of heterozygous *scospondin^stl297/+^* (J) and heterozygous *scospondin^stl300/+^* (K) demonstrating the C2262S and C879S non-synonymous mutations caused by each, respectively. (L) A schematic of the *Danio rerio* SCO-spondin protein domain structure.

**Figure S2.**
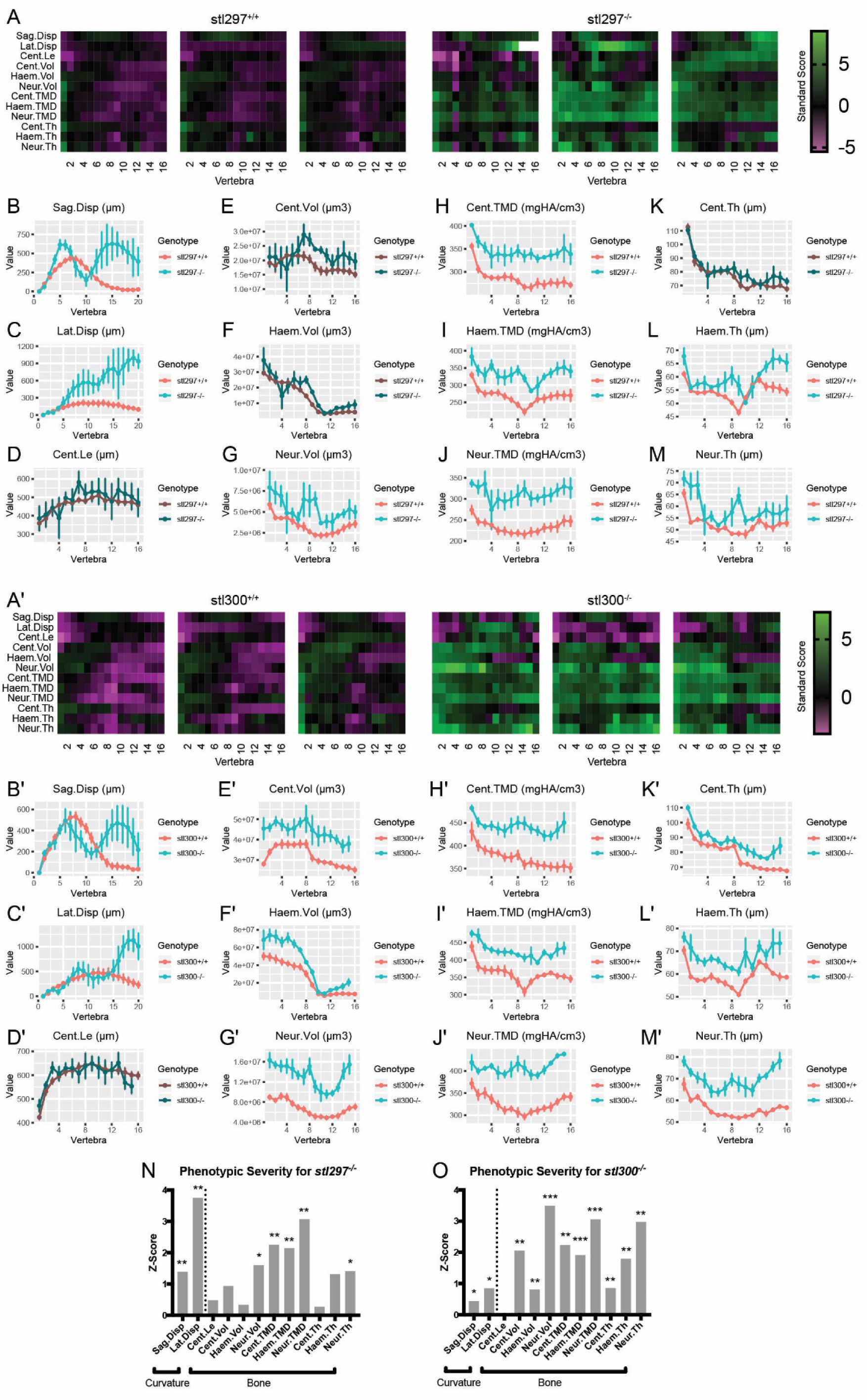
MicroCT analysis reveals increased spinal curvature, bone mass, and mineralization in *scospondin^stl297^* and *scospondin^stl300^* mutants. (A, A’) Heat maps for control and *scospondin^stl297^* (A) or *scospondin^stl300^* (A’) mutants. Each heat map represents a single fish (3 fish/group shown). Standard scores are computed as the difference between the value of the feature in the individual and the mean value of the feature across all vertebrae in the control population, divided by the standard deviation of the feature across all vertebrae in the control population. Note that for A, white boxes are those with standard scores >9. (B-M,B’-M’) Spinal curvature (B-C, B’-C’) and bone phenotypes (D-M, D’-M’) for *scospondin^stl297^* (B-M) or *scospondin^stl300^* (B’-M’) mutants. In each plot, the feature indicated by the graph heading (with units for y-axis) is plotted as a function of vertebra along the axial skeleton (mean ± SE). Plots associated with p<0.05 in the global test are colored in a lighter coloring scheme. For stl297 n=3-5/group (B-M), and for stl300 n=3-4/group (B’-M’). Sag.Disp, sagittal displacement; Lat.Disp, lateral displacement; Cent, centrum; Haem, haemal arch, Neur, neural arch; Vol, volume; TMD, tissue mineral density; Th, thickness; Le, length. (N, O) Average z-scores for data presented in A-M. P-values in the global test are indicated as follows: *: p<0.05, **: p<0.01, ***: p<0.001.

**Figure S3.**
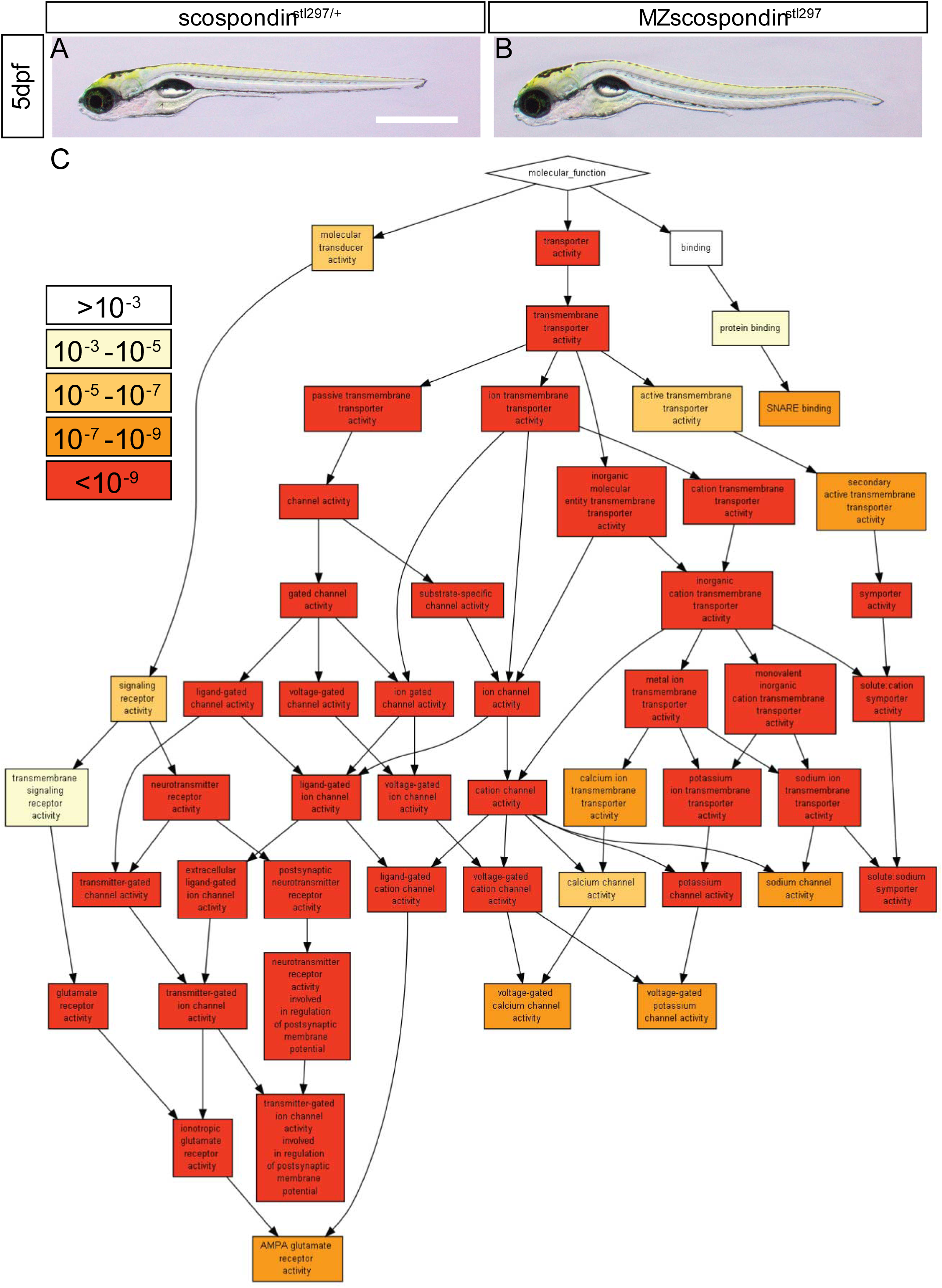
Mild notochord defects in 5dpf *MZscospondin^stl297^* and GO ontology ranking (p<1×10^-7^) of differentially expressed genes in *MZscospondin^stl297^*. (A-B) Brightfield lateral views of 5 dpf *scospondin^stl297/+^* (A) and *MZscospondin^stl297^* mutants (B). Scale bar is 1mm. (C) Gene ontology analysis was performed with the *GOrilla* gene ontology enrichment analysis and visualization tool, comparing *scospondin^stl297/+^ MZscospondin^stl297^* transcript levels at 5dpf.

**Figure S4.**
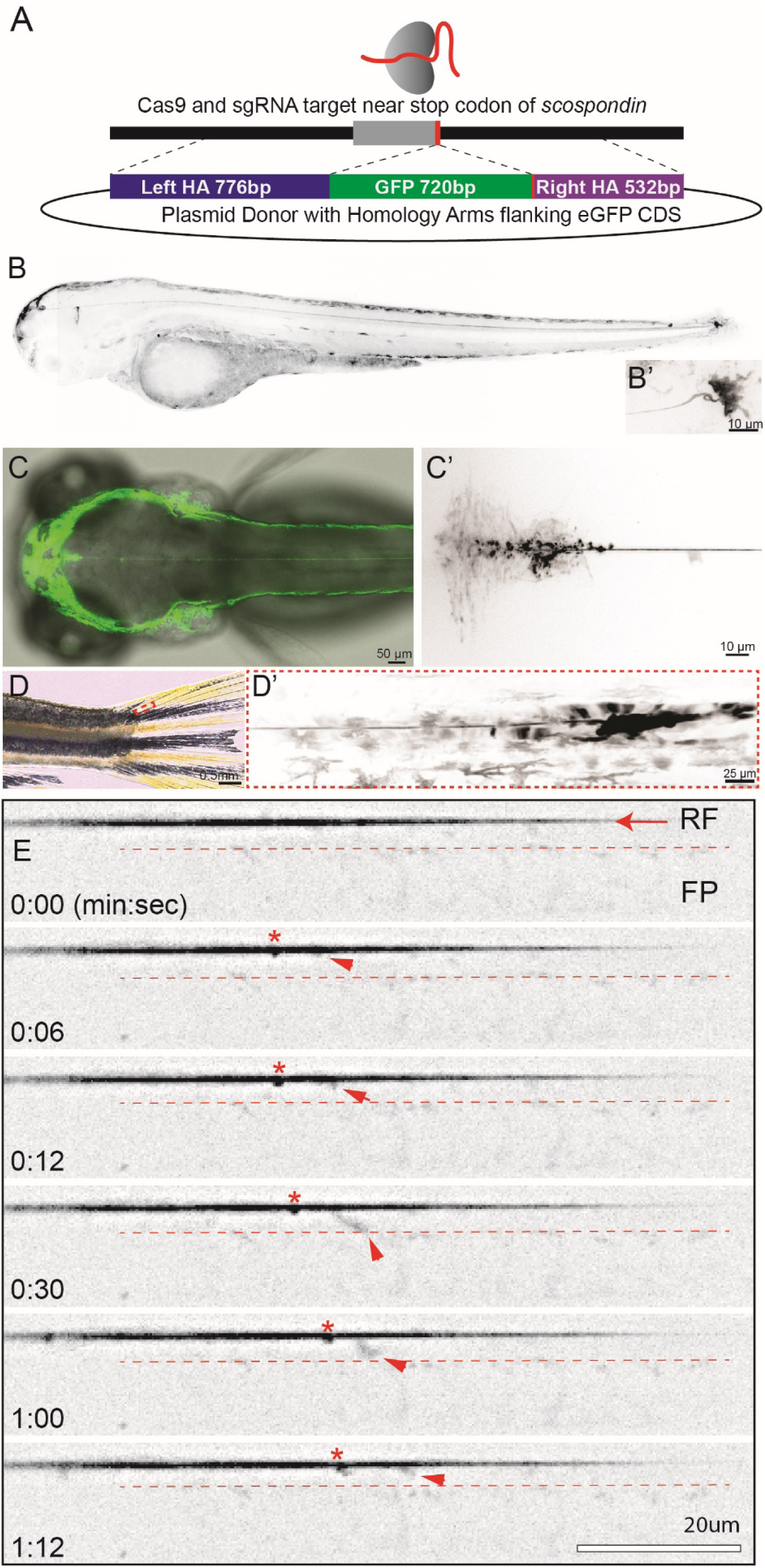
Description of engineering, fluorescent expression pattern, and Reissner fiber dynamics of the *scospondin-GFP^ut24^* knock-in zebrafish strain. (A) Schematic of CRISPR/Cas9 based genome editing with a donor plasmid containing EGFP coding sequence and homology arms. (B, B’) Inverted greyscale confocal image of a 3 dpf larval *scospondin-GFP^ut24^* knock-in line in a lateral view. (B’) High magnification image of the terminal ampulla region. (C, C’) DIC and confocal merge to highlight *scospondin-GFP^ut24^* expression and the Reissner fiber in a dorsal view of the head at 4 dpf, and (C’) an inverted greyscale image of *scospondin-GFP^ut24^* expression in the subcommissural organ. (D, D’) Bright field and confocal image of the *scospondin-GFP^ut24^* knock-in strain at 59 dpf. Inset is an inverted greyscale image of caudal most portion of the spinal canal to highlight the persistence of the Reissner fiber and terminal ampulla in juvenile fish. (E) Inverted greyscale confocal image of a 3 dpf larval *scospondin-GFP^ut24^* knock-in. RF = Reissner fiber labeled by a red arrow. FP = floor plate. Small red arrow points to processive puncta which appears to unravel and reach downward towards the floor plate. The red asterisks point at a punctum which moves along but is not observed to unravel until the last frame of the video.

**Video S1.** Confocal time-lapse inverted greyscale maximum intensity projection of *scospondin-GFP^ut24/+^* knock-in embryos during tailbud development (20-30 hpf) acquired at 5 min intervals. Time stamp is hr:min.

**Video S2.** Confocal time-lapse inverted greyscale maximum intensity projection of *scospondin-GFP^ut24/+^* knock-in embryos imaging the tail region during axial extension (2-3 dpf).

**Video S3.** Confocal time-lapse inverted greyscale maximum intensity projection of *scospondin-GFP^ut24/+^* knock-in embryos with photobleaching of the Reissner fiber (3 dpf).

**Video S4.** High-speed (10 Hz) confocal time-lapse inverted greyscale maximum intensity projection of the Reissner fiber in *scospondin-GFP^ut24/+^* knock-in embryos (3 dpf). Still images in Figure S4E.

**Table S1.** List of differentially expressed genes in 5dpf *MZscospondin^stl297^* mutants. Significantly altered genes were identified with DEseq2 analysis.

**Table S2:**
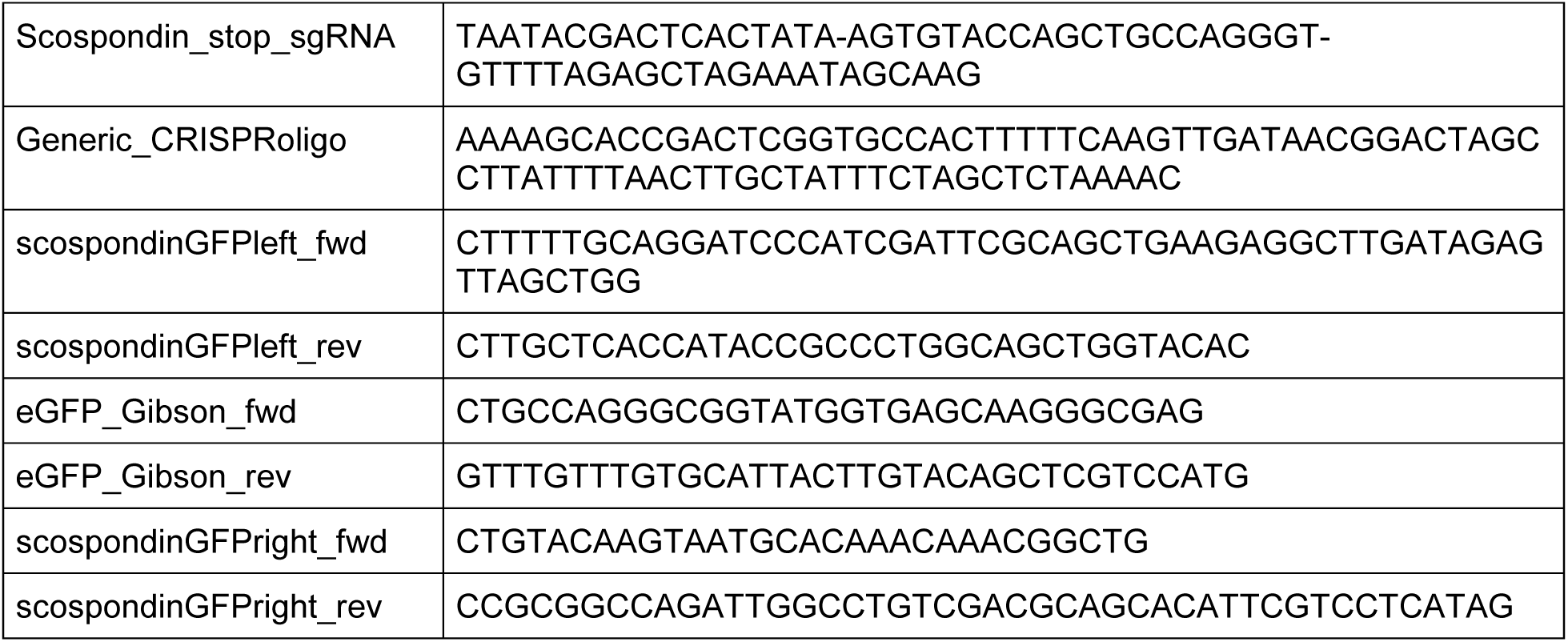
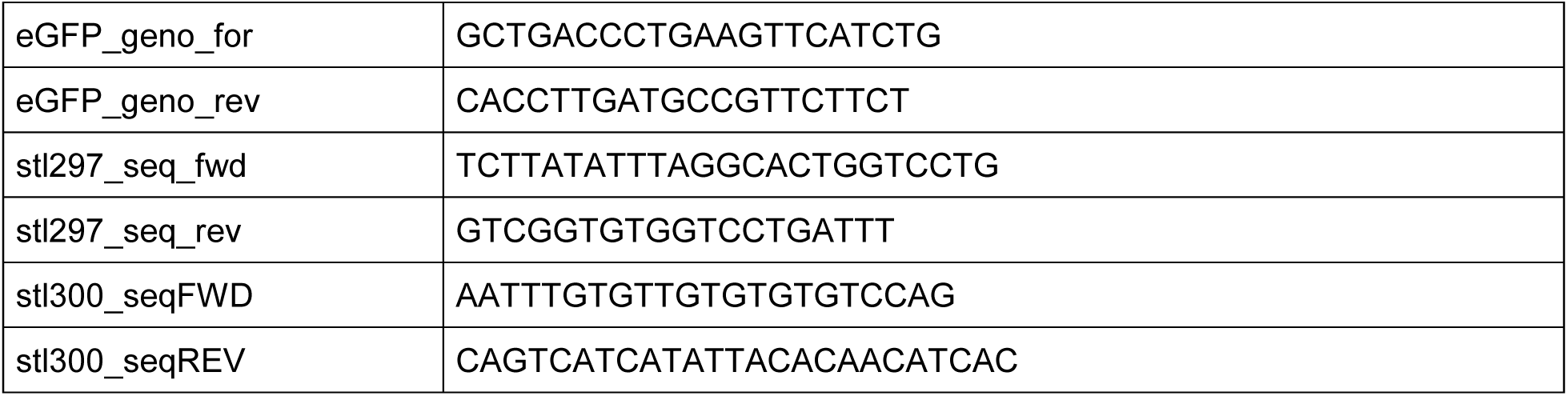
Oligos used in this manuscript.

KEY RESOURCES TABLE.

KEY RESOURCES TABLE
| REAGENT or RESOURCE | SOURCE | IDENTIFIER |
| --- | --- | --- |
| **Antibodies** |  |  |
| Rabbit anti-bovine Reissner fiber antiserum | Gift from Esteban Rodriguez and Maria Montserrat Guerra (Universidad Austral de Chile) | AFRU |
| Goat anti-Rabbit IgG (H+L) Alexa 488 | Invitrogen | A11034 |
| Goat anti-Rabbit IgG (H+L) Alexa 555 | Invitrogen | A21428 |
| Chicken anti-GFP | Abcam | ab13970 |
| Goat anti-Chicken IgY (H+L) Alexa 488 | Invitrogen | A11039 |
| **Chemicals, Peptides, and Recombinant Proteins** |  |  |
| EnGen Spy Cas9 NLS | New England Biolabs | M0646 |
| **Critical Commercial Assays** |  |  |
| CloneAmp Hif PCR Mastermix | Takara Bio | ST0506 |
| 5X In-Fusion HD Enzyme Premix | Takara Bio | ST0345 |
| HiScribe T7 High Yield RNA Synthesis Kit | New England Biolabs | E2040S |
| **Deposited Data** |  |  |
| RNAseq data | SRA browser | GSE138842 |
| WGS/WES data | SRA browser | GSE138920 |
| **Experimental Models: Organisms/Strains** |  |  |
| *scospondin*<sup>UT24</sup> | This manuscript |  |
| *scospondin*<sup>stl297</sup> | This manuscript |  |
| *scospondin*<sup>stl300</sup> | This manuscript |  |
| **Oligonucleotides** |  |  |
| See supplemental table S1 | This manuscript |  |
| **Recombinant DNA** |  |  |
| pCS108-scospondinGFP donor | This manuscript | NA |
| **Other** |  |  |
| Fiji | Schindelin et al., 2012 | https://fiji.sc/ |
| GraphPad Prism | Graph Pad Software | https://www.graphpad.com/scientific-software/prism/ |

## REFERENCES

1. Konjikusic, M.J., Yeetong, P., Boswell, C.W., Lee, C., Roberson, E.C., Ittiwut, R., Suphapeetiporn, K., Ciruna, B., Gurnett, C.A., Wallingford, J.B., et al. (2018). Mutations in Kinesin family member 6 reveal specific role in ependymal cell ciliogenesis and human neurological development. PLoS Genet 14, e1007817.

2. Grimes, D.T., Boswell, C.W., Morante, N.F., Henkelman, R.M., Burdine, R.D., and Ciruna, B. (2016). Zebrafish models of idiopathic scoliosis link cerebrospinal fluid flow defects to spine curvature. Science 352, 1341–1344.

3. Cantaut-Belarif, Y., Sternberg, J.R., Thouvenin, O., Wyart, C., and Bardet, P.L. (2018). The Reissner Fiber in the Cerebrospinal Fluid Controls Morphogenesis of the Body Axis. Curr Biol 28, 2479–2486 e2474.

4. Patten, S.A., Margaritte-Jeannin, P., Bernard, J.C., Alix, E., Labalme, A., Besson, A., Girard, S.L., Fendri, K., Fraisse, N., Biot, B., et al. (2015). Functional variants of POC5 identified in patients with idiopathic scoliosis. J Clin Invest 125, 1124–1128.

5. Hassan, A., Parent, S., Mathieu, H., Zaouter, C., Molidperee, S., Bagu, E.T., Barchi, S., Villemure, I., Patten, S.A., and Moldovan, F. (2019). Adolescent idiopathic scoliosis associated POC5 mutation impairs cell cycle, cilia length and centrosome protein interactions. PLoS One 14, e0213269.

6. Munoz, R.I., Kahne, T., Herrera, H., Rodriguez, S., Guerra, M.M., Vio, K., Hennig, R., Rapp, E., and Rodriguez, E. (2019). The subcommissural organ and the Reissner fiber: old friends revisited. Cell Tissue Res 375, 507–529.

7. Kohno, K. (1969). Electron microscopic studies on Reissner’s fiber and the ependymal cells in the spinal cord of the rat. Z Zellforsch Mikrosk Anat 94, 565–573.

8. Brand, M., Heisenberg, C.P., Warga, R.M., Pelegri, F., Karlstrom, R.O., Beuchle, D., Picker, A., Jiang, Y.J., Furutani-Seiki, M., van Eeden, F.J., et al. (1996). Mutations affecting development of the midline and general body shape during zebrafish embryogenesis. Development 123, 129–142.

9. Jaffe, K.M., Grimes, D.T., Schottenfeld-Roames, J., Werner, M.E., Ku, T.S., Kim, S.K., Pelliccia, J.L., Morante, N.F., Mitchell, B.J., and Burdine, R.D. (2016). c21orf59/kurly Controls Both Cilia Motility and Polarization. Cell Rep 14, 1841–1849.

10. Kramer-Zucker, A.G., Olale, F., Haycraft, C.J., Yoder, B.K., Schier, A.F., and Drummond, I.A. (2005). Cilia-driven fluid flow in the zebrafish pronephros, brain and Kupffer’s vesicle is required for normal organogenesis. Development 132, 1907–1921.

11. Meiniel, O., and Meiniel, A. (2007). The complex multidomain organization of SCO-spondin protein is highly conserved in mammals. Brain Res Rev 53, 321–327.

12. Meiniel, O., Meiniel, R., Lalloue, F., Didier, R., Jauberteau, M.O., Meiniel, A., and Petit, D. (2008). The lengthening of a giant protein: when, how, and why? J Mol Evol 66, 1–10.

13. Jeon, H., and Blacklow, S.C. (2005). Structure and physiologic function of the low-density lipoprotein receptor. Annu Rev Biochem 74, 535–562.

14. Rodriguez, E.M., Oksche, A., Hein, S., Rodriguez, S., and Yulis, R. (1984). Comparative immunocytochemical study of the subcommissural organ. Cell Tissue Res 237, 427–441.

15. Lehmann, C., and Naumann, W.W. (2005). Axon pathfinding and the floor plate factor Reissner’s substance in wildtype, cyclops and one-eyed pinhead mutants of Danio rerio. Brain Res Dev Brain Res 154, 1–14.

16. Eden, E., Navon, R., Steinfeld, I., Lipson, D., and Yakhini, Z. (2009). GOrilla: a tool for discovery and visualization of enriched GO terms in ranked gene lists. BMC Bioinformatics 10, 48.

17. Vera, A., Stanic, K., Montecinos, H., Torrejon, M., Marcellini, S., and Caprile, T. (2013). SCO-spondin from embryonic cerebrospinal fluid is required for neurogenesis during early brain development. Front Cell Neurosci 7, 80.

18. El-Bitar, F., Bamdad, M., Dastugue, B., and Meiniel, A. (2001). Effects of SCO-spondin thrombospondin type 1 repeats (TSR) in comparison to Reissner’s fiber material on the differentiation of the B104 neuroblastoma cell line. Cell Tissue Res 304, 361–369.

19. Kolmer, W. (1921). Das “Sagittalorgan” der Wirbeltiere. Zeitschrift für Anatomie und Entwicklungsgeschichte 60, 652–717.

20. Caprile, T., Hein, S., Rodriguez, S., Montecinos, H., and Rodriguez, E. (2003). Reissner fiber binds and transports away monoamines present in the cerebrospinal fluid. Brain Res Mol Brain Res 110, 177–192.

21. Buchan, J.G., Gray, R.S., Gansner, J.M., Alvarado, D.M., Burgert, L., Gitlin, J.D., Gurnett, C.A., and Goldsmith, M.I. (2014). Kinesin family member 6 (kif6) is necessary for spine development in zebrafish. Dev Dyn 243, 1646–1657.

22. Dendy, A. (1909). The Function of Reissner’s Fibre and the Ependymal Groove. Nature 82, 217–217.

23. Nicholls, G.E. (1917). Some Experiments on the Nature and Function of Reissner’s Fiber. The Journal of Comparative Neurobiology 27.

24. Hauser, R. (1969). [Dependence of normal tail regeneration in Xenopus larvae upon a diencephalic factor in the central canal]. Wilhelm Roux Arch Entwickl Mech Org 163, 221–247.

25. Rühle, H.-J. (1971). Anomalien im Wachstum der Achsenorgane nach experimenteller Ausschaltung des Komplexes Subcommissuralorgan—Reissnerscher Faden. Untersuchungen am Rippenmolch. Acta Zoologica 52, 23–68.

26. Thierer, J.H., Ekker, S.C., and Farber, S.A. (2019). The LipoGlo reporter system for sensitive and specific monitoring of atherogenic lipoproteins. Nat Commun 10, 3426.

27. Zappaterra, M.D., Lisgo, S.N., Lindsay, S., Gygi, S.P., Walsh, C.A., and Ballif, B.A. (2007). A comparative proteomic analysis of human and rat embryonic cerebrospinal fluid. J Proteome Res 6, 3537–3548.

28. Koch, M., Furtado, J.D., Falk, K., Leypoldt, F., Mukamal, K.J., and Jensen, M.K. (2017). Apolipoproteins and their subspecies in human cerebrospinal fluid and plasma. Alzheimers Dement (Amst) 6, 182–187.

29. Koch, S., Donarski, N., Goetze, K., Kreckel, M., Stuerenburg, H.J., Buhmann, C., and Beisiegel, U. (2001). Characterization of four lipoprotein classes in human cerebrospinal fluid. J Lipid Res 42, 1143–1151.

30. Parada, C., Escola-Gil, J.C., and Bueno, D. (2008). Low-density lipoproteins from embryonic cerebrospinal fluid are required for neural differentiation. J Neurosci Res 86, 2674–2684.

31. Farese, R.V., Jr., Ruland, S.L., Flynn, L.M., Stokowski, R.P., and Young, S.G. (1995). Knockout of the mouse apolipoprotein B gene results in embryonic lethality in homozygotes and protection against diet-induced hypercholesterolemia in heterozygotes. Proc Natl Acad Sci U S A 92, 1774–1778.

32. Vera, A., Recabal, A., Saldivia, N., Stanic, K., Torrejon, M., Montecinos, H., and Caprile, T. (2015). Interaction between SCO-spondin and low density lipoproteins from embryonic cerebrospinal fluid modulates their roles in early neurogenesis. Front Neuroanat 9, 72.

33. Li, H., and Durbin, R. (2009). Fast and accurate short read alignment with Burrows-Wheeler transform. Bioinformatics 25, 1754–1760.

34. Hur, M., Gistelinck, C.A., Huber, P., Lee, J., Thompson, M.H., Monstad-Rios, A.T., Watson, C.J., McMenamin, S.K., Willaert, A., Parichy, D.M., et al. (2017). MicroCT-based phenomics in the zebrafish skeleton reveals virtues of deep phenotyping in a distributed organ system. Elife 6.

35. Gistelinck, C., Kwon, R.Y., Malfait, F., Symoens, S., Harris, M.P., Henke, K., Hawkins, M.B., Fisher, S., Sips, P., Guillemyn, B., et al. (2018). Zebrafish type I collagen mutants faithfully recapitulate human type I collagenopathies. Proc Natl Acad Sci U S A 115, E8037–E8046.

36. Martin, M. (2011). Cutadapt removes adapter sequences from high-throughput sequencing reads. 2011 17, 3 % J EMBnet.journal.

37. Dobin, A., Davis, C.A., Schlesinger, F., Drenkow, J., Zaleski, C., Jha, S., Batut, P., Chaisson, M., and Gingeras, T.R. (2013). STAR: ultrafast universal RNA-seq aligner. Bioinformatics 29, 15–21.

38. Liao, Y., Smyth, G.K., and Shi, W. (2013). The Subread aligner: fast, accurate and scalable read mapping by seed-and-vote. Nucleic Acids Res 41, e108.

39. Love, M.I., Huber, W., and Anders, S. (2014). Moderated estimation of fold change and dispersion for RNA-seq data with DESeq2. Genome Biol 15, 550.

